# Brachyury controls *Ciona* notochord fate as part of a feedforward network and not as a unitary master regulator

**DOI:** 10.1101/2020.05.29.124024

**Authors:** Wendy M. Reeves, Kotaro Shimai, Konner M. Winkley, Michael T. Veeman

## Abstract

The notochord is a defining feature of the chordates. The transcription factor Brachyury (Bra) is a key regulator of notochord fate but here we show that it is not a unitary master regulator in the model chordate *Ciona*. Ectopic Bra expression only partially reprograms other cell types to a notochord-like transcriptional profile and a subset of notochord-enriched genes are unaffected by CRISPR Bra disruption. We identify Foxa.a and Mnx as potential co-regulators and find that combinatorial cocktails are more effective at reprograming other cell types than Bra alone. We reassess the network relationships between Bra, Foxa.a and other components of the notochord gene regulatory network and find that Foxa.a expression in the notochord is regulated by vegetal FGF signaling. It is a direct activator of Bra expression and has a binding motif that is significantly enriched in the regulatory regions of notochord-enriched genes. These and other results indicate that Bra and Foxa.a act together in a regulatory network dominated by positive feed-forward interactions, with neither being a classically-defined master regulator.

## Introduction

The notochord is the earliest chordate organ to form and plays essential structural and signaling roles in establishing the chordate body plan. The ascidian chordate *Ciona* has a particularly small and simple embryo in which invariant cell lineages give rise to a notochord of only 40 cells under the control of a compact ∼130Mb genome (Passamaneck and Di Gregorio, 2005; Satoh et al., 2003). As in all chordates, the *Ciona* notochord expresses the T-box transcription factor Bra (Corbo et al., 1997b) and loss of Bra function gives rise to major defects in notochord morphogenesis (Chiba et al., 2009; Satou et al., 2001; Yamada et al., 2003). Unlike in vertebrates, ascidian Bra is only expressed in the notochord and does not have broader roles in mesoderm induction (Corbo et al., 1997b; Yasuo and Satoh, 1993, 1998). In one of the earliest efforts to systematically define a tissue-specific transcriptional profile, Bra misexpression outside the notochord was used together with subtractive hybridization to identify 40 notochord-enriched genes induced directly or indirectly downstream of Bra (Hotta et al., 1999; Takahashi et al., 1999). Given that *Ciona* Bra is specifically expressed in the developing notochord, loss of function leads to notochord defects, and misexpression leads to the ectopic expression of a broad range of notochord target genes, it has widely been thought of as a canonical master regulator gene that is not only necessary but also sufficient for notochord fate.

There is strong evidence that Brachyury is a key regulatory of *Ciona* notochord fate, but the evidence that it acts as a unitary master regulator gene is weaker. Brachyury misexpression in the endodermal, A-line neural and notochord lineages under the control of the Foxa.a enhancer does not typically give rise to a larger notochord with additional intercalated cells, but instead to a mass of cells with little morphological differentiation (Takahashi et al., 1999). Several notochord enriched genes have been identified that are not induced by ectopic Bra expression (José-Edwards et al., 2011; Kugler et al., 2008), and several notochord enhancers have been identified that drive early notochord expression via essential Fox binding sites but with no recognizable Bra sites (José-Edwards et al., 2015).

We previously used RNAseq on flow-sorted *Ciona* notochord cells to identify a larger and more comprehensive set of ∼1300 transcripts enriched in the notochord during key stages of intercalation and elongation (Reeves et al., 2017). We also misexpressed Bra under the control of the Foxa.a enhancer similar to Takahashi et al, but using RNAseq as a more sensitive and quantitative readout. We identified over 900 upregulated genes, but found there was only modest overlap between the genes upregulated by ectopic Bra and the genes enriched in the notochord at comparable stages. This prompted the experiments described here, in which we use both gain-of-function and loss-of-function strategies coupled with modern transcriptional profiling to revise the gene regulatory network models for *Ciona* notochord fate specification. We find that Bra does not act as a unitary master regulator but instead as part of a densely interlocked positive feedforward network.

## Results

### Bra misexpression only induces a subset of notochord-enriched genes

Different tissues vary in their transcriptional and epigenetic states, so we misexpressed Bra in a broader range of cell types to determine if Bra could be more effective at inducing a notochord-like transcriptional profile in other contexts. In addition to the previously used Foxa.a enhancer, which drives expression in endoderm, A-line neural, notochord and mesenchyme cells (Reeves et al., 2017; Takahashi et al., 1999), we also used a pan-neural Etr1 (Celf3.a) enhancer (Sierro et al., 2006) and a muscle specific Tbx6-r.b enhancer (Kugler et al., 2010)(Fig 1A). Embryos were harvested for whole embryo RNAseq during notochord intercalation at early tailbud stage (Hotta stage 19) in triple biological replicates and gene expression was compared to control embryos electroporated with Bra>GFP. Reads were aligned using TopHat (Trapnell et al., 2012) and differential expression between conditions was tested by DESeq2 (Love et al., 2014) using a p-value threshold adjusted for multiple comparisons of 0.05.

**Figure 1.**
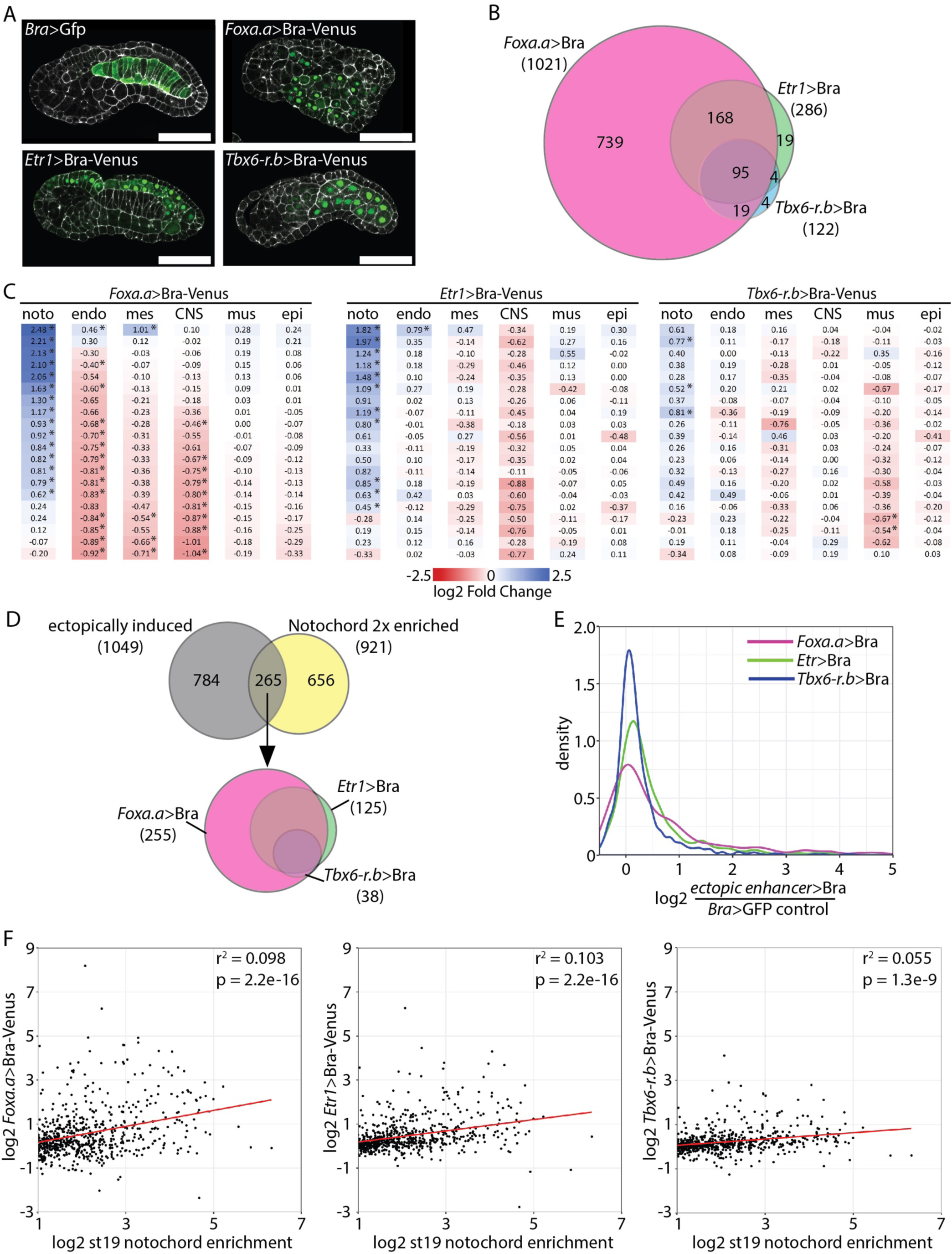
Ectopic expression of Bra upregulates a subset of notochord enriched genes. A) Venus-tagged Bra was misexpressed in *Ciona* embryos using the Foxa.a, Etr1 or Tbx6-r.b enhancers. Bra>GFP plasmid was used for control embryos. Phalloidin in white and Venus in green. Scale bar = 50 microns. B). Differential expression was calculated using DESeq. The overlap of all genes with significantly increased expression is shown. C). Effects of Bra misexpression on the expression of tissue specific markers. Marker genes highly enriched in notochord (noto), endoderm (endo), mesenchyme (mes), central nervous system (CNS), muscle (mus) and epidermis (epi) were identified using the scRNAseq data of (Cao et al., 2019). Log fold changes of Bra misexpression compared to control are shown. * indicates adjusted p-value is ≤ 0.05. D) Overlap of genes with at least two-fold enriched notochord expression (Reeves et al., 2017) and genes upregulated by Bra misexpression. E) Kernel density plot of the increase in expression by ectopic Bra for two-fold notochord enriched genes. F) Scatter plots of notochord enrichment at stage 19 in wildtype embryos (Reeves et al., 2017) versus the fold-change in response to Bra misexpression driven by the different constructs. Only genes with a statistically significant stage 19 notochord enrichment of at least two-fold are shown. The red lines show the best fit linear regressions.

We found that Brachyury misexpression using the neural and muscle drivers led to considerably fewer genes being identified as upregulated (Foxa.a>Bra: 1021; Etr1>Bra: 286; Tbx6-r.b >Bra: 122). These ectopic expression constructs induced largely overlapping sets of genes (Figure 1B, Supp Table1), with the genes upregulated by the muscle and neural drivers representing a small subset of the genes induced by Foxa.a>Bra.

If Bra is a master regulator of notochord fate, then misexpression outside the notochord should lead to a downregulation of tissue specific markers in the ectopically expressing tissues and an upregulation of notochord-enriched genes. To test this, we used recent *Ciona* single cell RNAseq data (Cao et al., 2019) to identify the twenty genes that are most differentially expressed in each target tissue at stage 19 (Supp Table 2). Although many target tissue markers were downregulated to some extent, the fold changes were quite variable. Many showed minimal change, and of those that went down many were not statistically significant. A few were paradoxically upregulated. (Figure 1C). Notochord marker genes tended to be upregulated, but again many showed little or no effect. This suggests that, unlike with a classically defined master regulator gene, ectopic misexpression of Bra can only induce a partial reprogramming of other tissues.

When we looked at the full set of notochord enriched genes previously identified by FACS-seq (Reeves et al., 2017), relatively few were upregulated by ectopic Bra expression. Only 293 of the 1372 genes (21%) that are significantly notochord enriched at either stage 16 and/or stage 19 were statistically significantly upregulated by any of the ectopic Bra misexpression constructs (Supp Table 3). The 1372 notochord-enriched genes span a broad range of enrichment levels, including many with expression that is only slightly higher in the notochord compared to other cell types. If we restrict this comparison to genes that are at least two fold enriched in notochord, then 265 of the 921 twofold-enriched genes (29%) were ectopically induced (Figure 1D,E), but the majority showed no statistically significant change. There is a relationship between strength of notochord enrichment and ectopic upregulation by Bra, but there were still many highly enriched notochord genes that were not upregulated by ectopic Bra expression (Figure 1F and Supp Table 3). This correlation between notochord enrichment and the degree of ectopic upregulation by Bra was statistically significant for all three constructs, but the correlation coefficients were very low (0.06-0.1).

Bra misexpression in diverse tissues can upregulate many notochord genes, but it is not sufficient to upregulate expression of all notochord enriched genes, including some of the most highly enriched.

### Some notochord-enriched genes are independent of Brachyury

Even if Brachyury expression is not sufficient to completely transform other cell types to notochord, it could still be a unitary regulator alone at the top of the transcriptional cascade for notochord differentiation. To address the necessity of Bra function in the expression of notochord-enriched genes, we used CRISPR-Bra RNP egg injections to disrupt Bra and examined the effect on gene expression at early neurula stage (Hotta stage 14). A germline Bra mutation exists, but the CRISPR method allowed us to quantify early changes in gene expression before homozygous mutant embryos can reliably be identified. We used a targeting RNA specific to the third exon of Bra, capable of replicating the tail elongation defect of homozygous mutants (Chiba et al., 2009)(Figure 2A). TIDE analysis (Brinkman et al., 2014) confirmed a somatic mutation efficiency ranging from 67% to 77% in the three biological replicates harvested for RNAseq (Fig 2B). It was impractical to flow-sort notochord cells from injected embryos, so we used RNA from whole embryos. Assuming that Brachyury inhibition will only affect notochord expression and not expression elsewhere in the embryo, the maximum possible fold decrease for a given gene is a function of its notochord enrichment levels, which we have previously estimated by FACS-seq (Reeves et al., 2017). The red line in Fig 2C indicates the theoretical maximum decrease across the range of St16 notochord enrichment levels, as described in methods.

**Figure 2.**
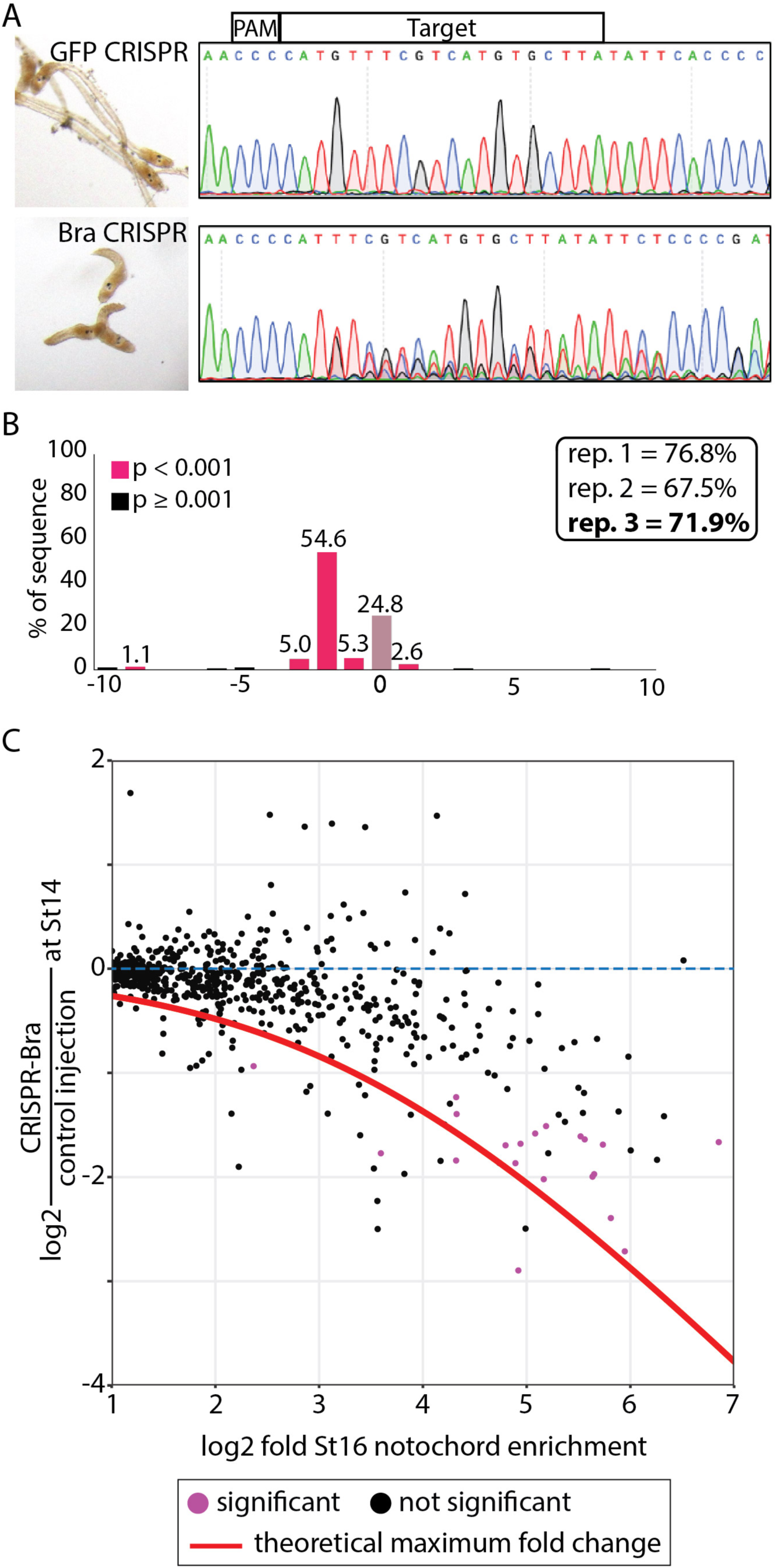
Somatic CRISPR disruption of Bra. A) Eggs were injected with RNP-guide RNA complexes targeting either GFP (control, top) or Bra (bottom). Representative embryos at late tailbud are shown on the left with sequencing results used for TIDE analysis of the Bra locus on the right. B) TIDE results from one replicate, showing efficient mutagenesis giving rise largely to a two base pair deletion in injected embryos. TIDE-estimated indel rates for the three RNAseq replicates are shown in the inset. C) Notochord enrichment at stage 16 (Reeves et al., 2017) compared to change in expression at stage 14 in CRISPR-Bra injected embryos versus controls. Only genes with a statistically significant stage 16 notochord enrichment of at least two-fold are shown. Red line indicates the theoretical maximum decrease observable in whole embryo RNAseq for a given notochord enrichment value.

As expected, expression of notochord-enriched genes tended to be decreased, with genes that are more notochord-enriched showing larger responses (Fig 2C). There was, however, a broad range of fold changes, with some highly notochord-enriched genes showing little or no response to Bra disruption, including Salla, Col2a1, Col11A1/2 and CHDH. To validate the RNAseq results, we selected several notochord genes spanning a range of notochord enrichment values and fold changes in response to Bra CRISPR to test by in situ hybridization in Bra crispant and control embryos (Fig 3). As expected, notochord expression was lost for strongly notochord-enriched genes showing large decreases in the Bra CRISPR RNAseq. We included a Brachyury probe as a control in these experiments and found that the loss of Bra expression was similar in penetrance to the loss of these highly Bra-dependent transcripts. A similar loss of notochord expression was also seen for several moderately notochord-enriched genes with smaller responses to Bra CRISPR. Normal notochord expression was confirmed, however, for several highly notochord-enriched genes showing minimal changes in response to Bra CRISPR, including the notochord-enriched transcription factor Foxa.a. This indicates that some aspects of notochord-enriched gene expression are independent of Bra.

**Figure 3.**
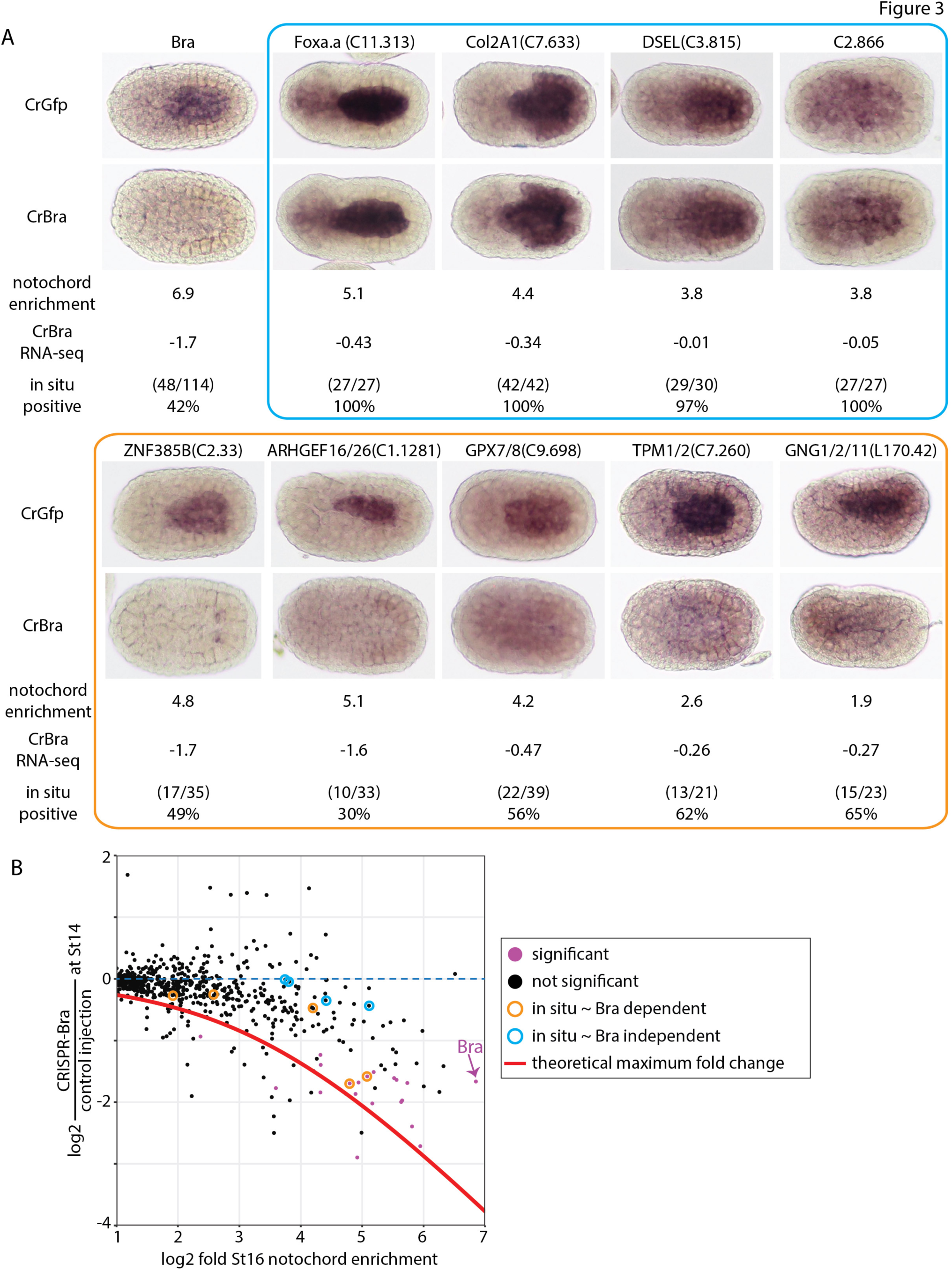
Expression patterns of notochord-enriched genes in Bra crispant embryos. A) Injected embryos were fixed at stage 16 and stained by in situ hybridization with probes for the indicated genes. Since mosaic expression was not observed in control embryos, Bra crispant embryos were scored as positive if they had uniform expression in the notochord and negative if there was either a total loss of notochord expression or a mosaic loss in a subset of notochord cells. Representative embryos are shown along with log2 notochord enrichment at Stage 16 (Reeves et al., 2017) and log2 fold change in Bra crispant embryos. The blue and orange boxes indicate nominally Bra-independent genes and Bra-dependent genes. B) Scatter plot of notochord enrichment and Bra CRISPR fold change comparable to Fig. 2C, but now highlighting the specific genes tested by in situ hybridization.

### Foxa.a and Mnx are candidate factors to act in parallel with Bra

Given that Bra is neither necessary in CRISPR loss of function assays nor sufficient in ectopic expression assays for the expression of all notochord-enriched genes, there are likely to be additional transcription factors working in parallel to Bra at the top of the notochord GRN. Our inferences based on genome-wide transcriptional profiling are also supported by prior observations. Kugler et al (Kugler et al., 2008) identified six genes expressed in notochord after gastrulation that were neither induced by Foxa.a>Bra misexpression nor repressed by expression of a Bra-Engrailed repressor fusion protein. The LMX-like transcription factor has also been shown to retain some notochord expression in Bra homozygous mutant embryos (José-Edwards et al., 2011).

In our misexpression studies, the extent of notochord gene upregulation varied greatly between different target tissues, with the Foxa.a>Bra construct having the greatest effect (Fig 1D). One possibility is that these tissues are uniquely sensitive to Bra misexpression because Foxa.a is a coregulator with Bra of notochord fate. Foxa.a is required for notochord fate, but it is difficult to assess whether it acts at least in part in parallel to Brachyury because it is also an upstream regulator of Bra expression (Imai et al., 2006; Kumano et al., 2006). Foxa.a is an essential and direct upstream regulator of ZicL (Imai et al., 2002b), and ZicL is an essential and direct upstream regulator of Bra (Imai et al., 2002b; Yagi et al., 2004). Important FoxA sites have been found in several notochord enhancers (José-Edwards et al., 2015; Passamaneck et al., 2009), but Foxa.a’s specific roles after the induction of ZicL at the 32 cell stage are not well understood. Given its strong, Bra-independent notochord expression in neurula stage embryos, it could easily be acting in parallel to Bra as well as upstream.

Notochord fate and Brachyury expression are dependent on vegetal FGF signaling (Imai et al., 2006, 2002a; Yasuo and Hudson, 2007), so we hypothesized that other parallel regulators of notochord fate might share this same dependence. To identify such genes, we inhibited FGF signaling with the MEK inhibitor U0126 at the 32 cell stage and then performed RNAseq at the 64 and 112-cell stages to catch the earliest changes in gene expression between drug-treated and control embryos. A total of 152 genes had significantly decreased expression at either stage (14 at the 64 cell stage and 150 at the 112 cell stage), including 16 transcription factors (Fig 4C, Supp Table 4). We compared these results with published in situ hybridization and single cell RNAseq data and found that 4 of the 16 are specifically expressed in other FGF-dependent tissues, like endoderm, nerve cord or mesenchyme, and thus are not likely involved in notochord specification. Nine transcription factors were expressed broadly in the endoderm, nervous system, mesenchyme and notochord of the early embryo. Of the remaining genes, Foxa.a was slightly but significantly decreased (p-value < 0.1) at the 112-cell stage in U0126 treated embryos and both Bra and Mnx were strongly and significantly down at both stages (p-value < 0.05). During gastrulation, Mnx is expressed in medial row I of the neural plate and primary muscle (Hudson et al., 2007), and becomes specific to the visceral ganglion after neurulation (Imai et al., 2009, 2004). However, it also has specific but transient expression in the notochord as it becomes fate-restricted at the 64-112 cell stages (Imai et al., 2006, 2004). Mnx is a homeodomain transcription factor previously hypothesized to play a role in repressing neural genes in the early notochord (Kodama et al., 2016).

**Figure 4.**
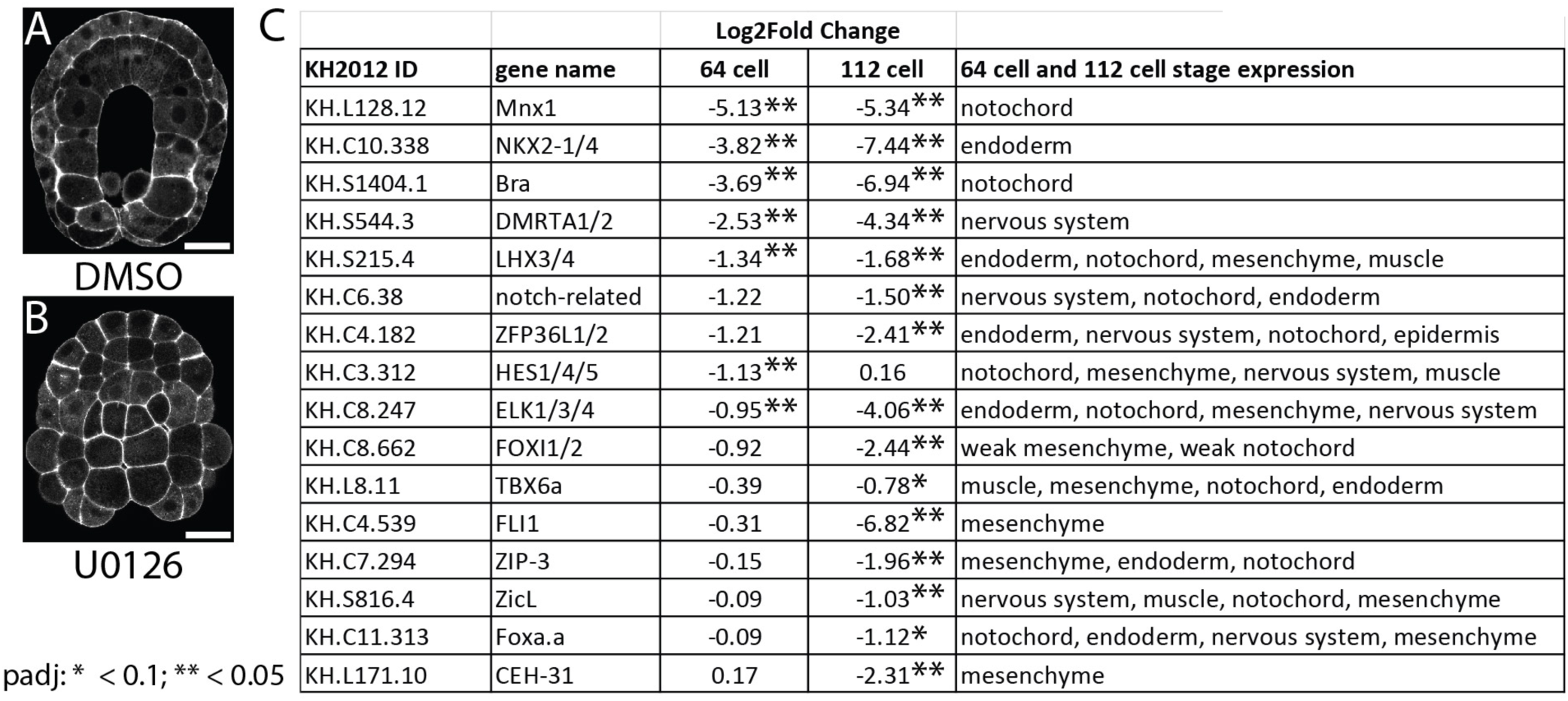
Transcription factors dependent on early FGF signaling. A,B) Embryos were treated with A) DMSO or B) U0126 at the 32 cell stage then harvested for RNAseq at the 64 (not shown) and 112 cell stages. Scale bar = 50 µm. C) Transcription factors with decreased expression in response to inhibition of FGF signaling. The expression patterns noted are based upon publicly available in situ data from ANISEED (Dardaillon et al., 2020) and/or single cell RNAseq (Cao et al., 2019). See supplemental table 4 for more details.

### Combined cocktails are more effective at reprogramming neural plate to notochord

Foxa.a and Mnx are both strong candidates for transcription factors that could function in parallel with Bra at the top of the notochord GRN. We used the neural enhancer Etr1 to express Mnx and/or Foxa.a in combination with Bra (Fig 5A) to determine if combinatorial expression of these transcription factors was more efficacious than Bra alone in upregulating notochord enriched genes. We chose this tissue specific enhancer because Etr1>Bra misexpression had an intermediate effect on gene expression profiles compared to Foxa.a>Bra and Tbx6-r.b >Bra. Etr1 expressing neural tissues were less refractory to reprogramming than muscle (Fig1), and only partially overlap with the endogenous expression of Foxa.a. Whole embryo expression at stage 19 was measured by RNAseq and compared to expression in embryos electroporated with Etr1>H2B-Venus (Supp Table 5).

**Figure 5.**
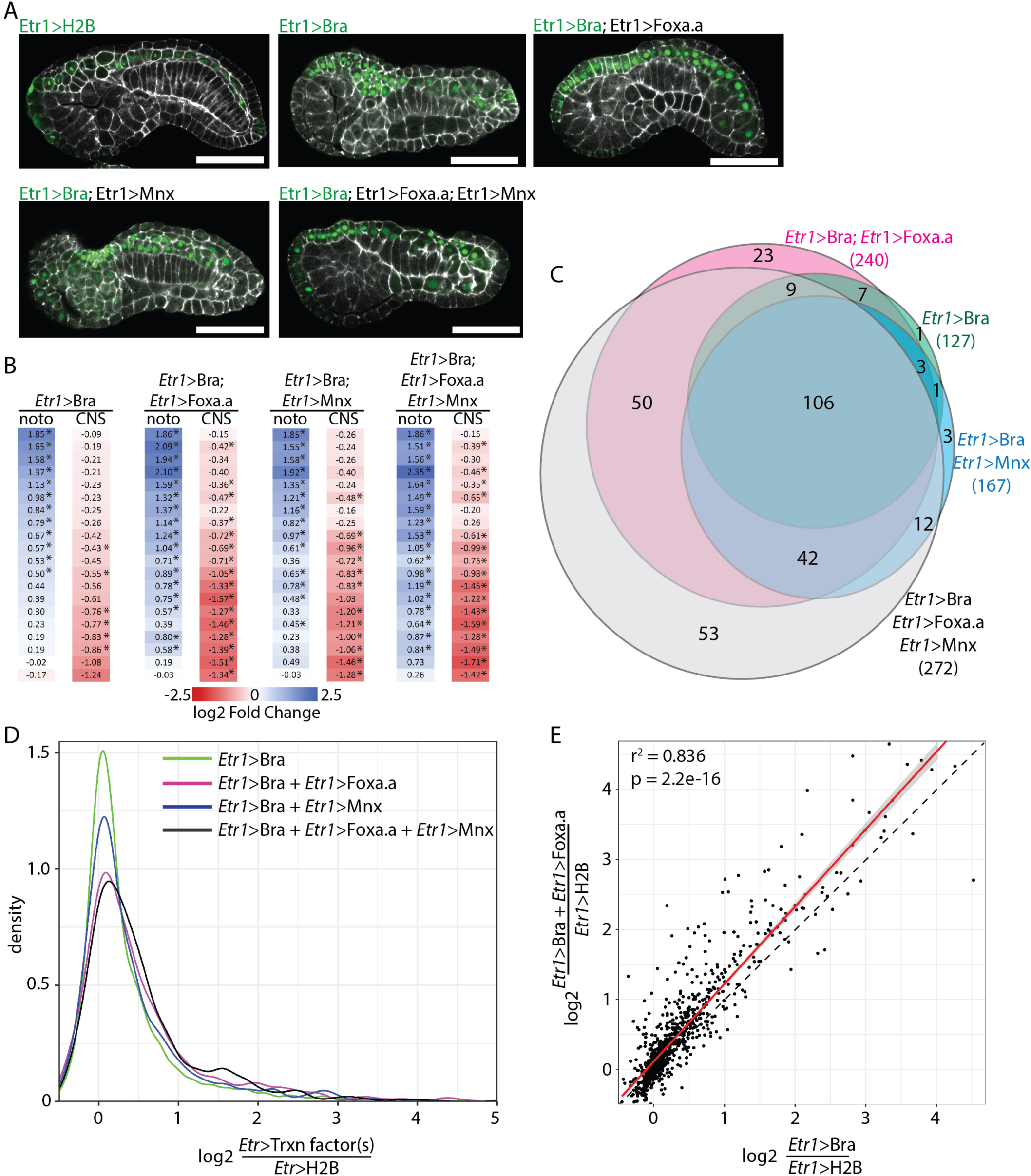
Combinatorial misexpression of Bra, Foxa.a and Mnx. A) Representative confocal images of embryos expressing different combinations of Bra-Venus, Foxa.a-myc and/or Mnx-HA fusion proteins under the control of the pan-neural Etr1 enhancer. Etr1>H2B-Venus was expressed in control embryos. Embryos were collected for RNAseq and confocal imaging at stage 19. Phalloidin is shown in white and Venus in green. Scale bar = 50 microns. B) Effect of transcription factor misexpression on tissue specific markers derived from (Cao et al., 2019) as in Figure1. noto = notochord and CNS = central nervous system. * = adjusted p-value ≤ 0.05. C) Overlap of notochord two-fold enriched genes whose expression is significantly upregulated by misexpression of indicated transcription factors. D) Kernel density plot of the fold-change of expression for two-fold enriched notochord genes in response to different ectopic transcription factor combinations. E) Comparison of differential expression due to Bra misexpression alone vs the Bra/Foxa.a combination. Best fit linear regression shown in red flanked by the *95%* confidence interval. The dotted line indicates a perfect 1:1 relationship.

As predicted, co-expression of Foxa.a and/or Mnx with Bra had a greater effect on the induction of notochord markers and repression of neural markers than Bra alone (Fig 5B, Supp Table 2), although some markers still remained largely unchanged.

We next looked at the 921 genes that are enriched at least 2-fold in notochord according to our earlier FACS-RNAseq experiment (Reeves et al., 2017). Co-expression of Mnx and Bra had a significant effect on only a few notochord enriched genes (Fig 5C,D, Supp Table 3), suggesting a more limited or gene-specific role for Mnx in the regulation of notochord gene expression. However, since Mnx is normally expressed in notochord for only a short window around the 64/112 cell stages (Imai et al., 2006, 2004), it is possible that a short pulse of misexpression might have different effects than the protracted misexpression performed here. Additional work will be necessary to fully test the potential roles of Mnx in notochord fate induction.

In contrast, co-expression of Foxa.a and Bra had a larger effect on the upregulation of notochord enriched genes. Bra/Foxa.a misexpressing embryos had almost twice as many notochord enriched genes upregulated compared to Bra alone (240 vs 127; 26% of 2x notochord enriched genes vs 14%) (Fig 5C, D). Their co-expression also increased the expression of most genes already upregulated by Etr1>Bra alone (Fig 5E).

We saw little synergy when simultaneously misexpressing all three genes. The triple Bra/Foxa.a/Mnx cocktail gave similar effects to Bra/Foxa.a, but with the addition of a few genes specifically affected by Mnx expression. Simultaneous mixexpression of Bra and Foxa.a in the neural plate did not upregulate all notochord-enriched genes, but was considerably more potent than Bra alone. These results support the hypothesis that Foxa.a is not just an indirect upstream activator of Bra expression, but also acts in parallel to Bra to regulate notochord target genes. Mnx’s role is not clear, but it is less important than Foxa.a in misexpression assays. FoxA is known to be a key regulator of notochord fate in both vertebrates (Ang and Rossant, 1994; Weinstein et al., 1994) and tunicates (Imai et al., 2006; José-Edwards et al., 2015; Kumano et al., 2006), but the details of the regulatory networks connecting FoxA, Bra and downstream targets are not well understood in any model. We pursued several parallel strategies to clarify these relationships.

### Notochord Foxa.a expression depends on vegetal FGF signaling

Foxa.a is a major determinant of anterior vegetal fates, with a complex expression pattern in A-line neural, notochord and endodermal cells. Foxa.a expression is initially induced throughout the vegetal region by beta-catenin signaling at the 16 cell stage (Imai et al., 2006). By early gastrulation, it is expressed most strongly in notochord, a pattern that continues through neurulation (Corbo et al., 1997a; Di Gregorio et al., 2001) and early tailbud stages (Imai et al., 2004). In contrast, early CNS expression of Foxa.a is lost by late gastrula stage, and then turned back on in only the ventral-most neuroepithelium during neurulation (Corbo et al., 1997a). Expression in the endodermal strand is weak but stable throughout embryonic development (Corbo et al., 1997a; Di Gregorio et al., 2001). Inhibition of FGF signaling by a MEK inhibitor at the 32 cell stage results in a weak but significant decrease in Foxa.a expression by whole embryo RNAseq at the 112 cell stage (see above/Fig 4). Although FGF dependence was not previously seen in *Ciona* by whole embryo RT-PCR (Imai et al., 2006), notochord expression of FoxA in *Halocynthia* was reduced at the 64 cell, but not 32 cell, stage by injection of a FGF9/16/20 morpholino (Kumano et al., 2006). This raises the question of whether its later notochord expression in *Ciona* is comparable to Bra in being dependent on the vegetal FGF signals at the 32 cell stage that are required for notochord fate induction. We treated embryos with U0126 at the 32-cell stage and then cytochalasinB at the 112 cell stage, to arrest cell division and allow for clear identification of cell types in older embryos. Major cell fates including notochord are known to be established correctly in cleavage-arrested embryos (Crowther and Whittaker, 1986; Whittaker and Meedel, 1989) and we have previously shown that cytB treatment does not have a significant effect on expression of notochord-enriched genes (Reeves et al., 2017). At stage 16 (mid notochord intercalation in untreated embyros), Foxa.a expression has been lost specifically in the notochord but not in the presumptive endoderm (Fig 6A). This indicates that notochord-enriched Foxa.a expression in *Ciona* has essential direct or indirect inputs from FGF signaling specifically in the notochord lineage and distinct from its FGF-independent regulation at early cleavage stages. We have not mapped out precise start or end points to this dependency on FGF, but it identifies an important new edge in the *Ciona* notochord GRN.

**Figure 6.**
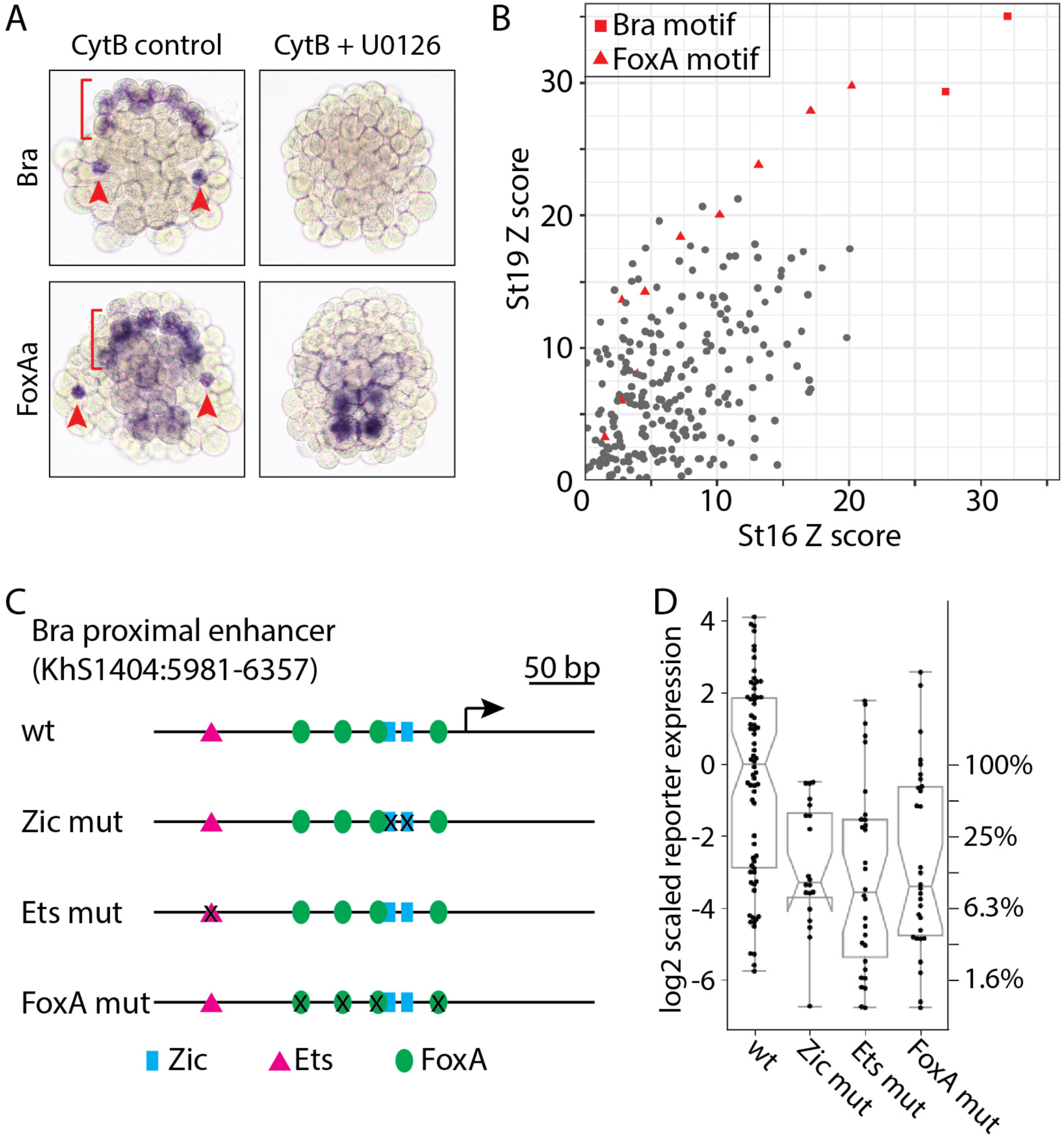
Cis-regulatory interactions involving Foxa.a and Bra. A) In situ hybridization for Bra and Foxa.a with and without inhibition of early FGF signaling. Embryos were treated at 32 cell stage with the MEK inhibitor U0126 or DMSO vehicle control, then cleavage arrested at the 112 cell stage with Cytochalasin B before fixing at a timepoint equivalent to Stage 16. Red arrowheads indicate secondary notochord and red bracket indicates the arc of primary notochord. B) Transcription factor binding site enrichment analysis for putative regulatory regions of genes enriched at least two fold in notochord at stage 16 or stage 19 compared to the putative regulatory regions of genes depleted in the notochord. Z-scores for all enriched TFBSs are shown, with Bra and FoxA motifs highlighted in red. C,D) Quantitative reporter assay analysis of specific predicted transcription factor binding site mutations in the proximal Bra enhancer region. C) shows a map of the Brachyury upstream region with predicted binding sites for Ets (magenta triangle), Zic (blue box) and FoxA (green oval) labeled. D) Reporter expression for the different TFBS mutation constructs. Each point represents a different embryo imaged by 3D confocal microscopy with notochord expression of the reporter quantified as described in the Methods. This confocal-based assay has high dynamic range and there is considerable embryo to embryo variation due to transgene mosaicism. The reporter activity scores are normalized to give the control construct a log-transformed median of zero. A separate y-axis is used on the right to show linear-scaled percent decreases with respect to the median of the control plasmid. The bar and whisker plots show the medians, quartiles and minima/maxima.

### Foxa.a and Bra sites are both enriched near *Ciona* notochord genes

An alternate strategy for inferring cell-type specific transcriptional regulators is to identify transcription factor binding motifs enriched in the putative regulatory regions of genes with tissue-specific expression patterns. If Foxa.a is an important co-regulator of notochord gene expression acting not just upstream of Bra but also in parallel, then FoxA binding motifs should be overrepresented in the enhancers of notochord enriched genes. (José-Edwards et al., 2015) previously found that several CRMs capable of driving notochord expression of a reporter at mid-tailbud frequently contained functionally important Bra and Fox binding sites, but did not address whether these or other binding motifs are statistically enriched across notochord genes overall. Transcription factor binding site (TFBS) enrichment analysis also provides an alternate means of predicting regulators that may have been overlooked based on complex or imperfectly annotated expression patterns.

We focused again on genes enriched at least two-fold in notochord during neurulation (stage 16) as well as early tailbud (stage 19). *Ciona* enhancers tend to be near to the transcriptional start site (Irvine, 2013), so we used a window 1500 bp upstream and downstream from the transcription start site of each gene model from which we subtracted predicted exons (see methods for full details). Putative regulatory regions of notochord enriched genes were tested for overrepresented TFBSs using the 2018 JASPAR vertebrate TFBS profile set (Khan et al., 2018) and the oPOSSUM 3.0 Single-Site Analysis algorithm (Kwon et al., 2012), with the enhancers from the 1000 most notochord depleted genes at each stage as the control set. As expected, T-box binding sites predicted to be bound by Bra had the highest Z-scores reflecting the number of TFBS occurrences in the notochord vs control set. FoxA binding sites were the next most significantly overrepresented motif in the notochord enriched set at both stages (Figure 6B), consistent with a key role for Foxa.a in regulating notochord gene expression. We also saw modest enrichment for several other motifs whose associated transcription factors have putative roles in notochord differentiation including TBX2 (José-Edwards et al., 2013), Fos/Jun related bZIPs(José-Edwards et al., 2011), and some homeodomain motifs (José-Edwards et al., 2011; Katikala et al., 2013), but Bra and FoxA sites were by far the most enriched. Mnx is a homeodomain TF with a very short and common consensus motif, so enrichment would not necessarily be expected.

### Foxa.a is a direct activator of *Ciona* Brachyury expression

Foxa.a acts indirectly upstream of Brachyury via induction of ZicL, and also acts in parallel to Brachyury in the induction of downstream notochord genes, but it is also possible that it could be acting directly on Brachyury as well. This is supported by work in *Halocynthia* suggesting that FoxA and Zic act at least partly in parallel as competence factors for induction of Bra expression by FGF (Kumano et al., 2006). Potential FoxA binding sites have been identified in a region of the *Halocynthia* Bra enhancer, but their mutation did not decrease Bra>reporter expression in a LacZ reporter assay (Matsumoto et al., 2007). Potential roles for Foxa.a sites in the cis-regulatory control of *Ciona* Bra have not previously been tested. To determine if Foxa.a is a direct activator of *Ciona* Bra expression, we quantified the effects of mutations in Ets, Zic and FoxA binding sites in the 377bp Bra proximal enhancer (KhS1404:5981-6357) using a sensitive, high-dynamic range assay based on confocal microscopy. Ets and Zic were used as positive controls as they have previously been shown to be essential and direct positive regulators of Bra expression (Farley et al., 2016; Matsumoto et al., 2007; Yagi et al., 2004). Wildtype and mutant enhancers were cloned upstream of a basal pFOG promoter and Venus reporter (Harder et al., 2019; Stolfi et al., 2015) and electroporated into embryos in triple biological replicates (Fig 6C). At least 10 embryos were imaged for each construct in each biological replicate and reporter expression was quantified using a method similar to (Harder et al., 2019). As seen previously, Zic and Ets sites were critical for wt expression, with mutations reducing median expression down to 8-10% of that seen with a wildtype reporter (Fig 6D). Mutation of 4 FoxA sites within the enhancer similarly reduced median expression to 9.5% of wildtype levels, supporting the hypothesis that Bra expression is directly Foxa.a dependent.

### Feedforward loops in the *Ciona* notochord GRN

While we did not identify any entirely new nodes (trans-acting factors) in the *Ciona* notochord GRN, we identified multiple new edges (regulatory interactions) (Fig 7). These demonstrate that notochord fate specification is dominated by interlocked coherent feed-forward interactions. FGF is not only a direct activator of Brachyury, but also an activator of Foxa.a. Foxa.a is not only an indirect activator of Bra via ZicL but also a direct activator. Notochord target genes are induced not only by Bra but also by other factors including Foxa.a. All of the major regulators of notochord fate specification act positively at multiple hierarchical levels.

**Figure 7.**
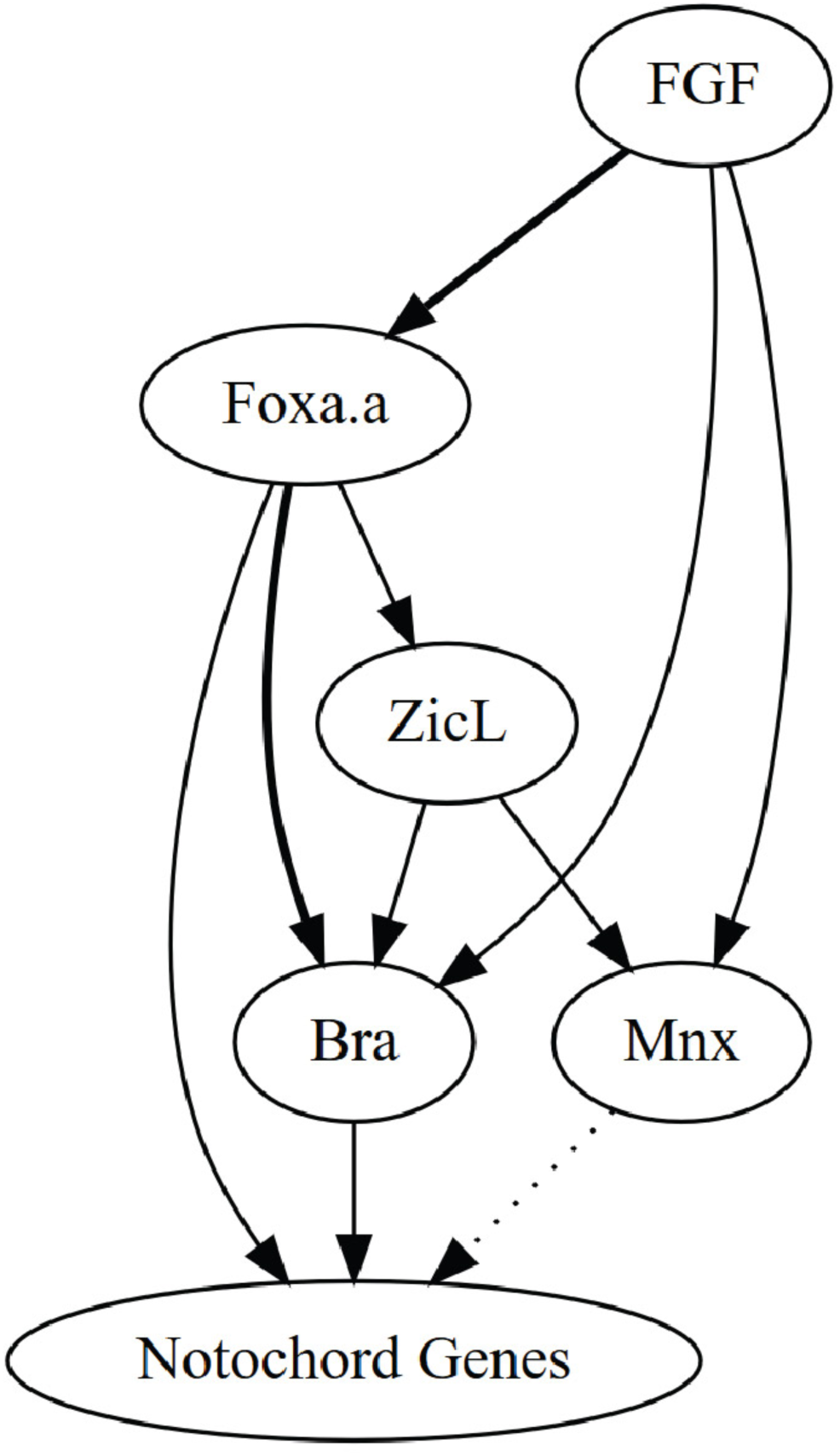
A GRN for *Ciona* notochord fate specification. Bold lines indicate new cis-regulatory relationships in the *Ciona* notochord GRN identified or validated in this paper. The dotted arrow from Mnx is supported by overexpression results but remains to be confirmed by loss of function. Other regulatory interactions in the *Ciona* GRN were previously described in (Imai et al., 2006, 2002a, 2002b; Yagi et al., 2004; Yasuo and Hudson, 2007).

## Discussion

Coherent feed-forward loops where A induces B and then A and B jointly induce C are common in biological networks. Comparable loops have been identified in diverse models of cell fate specification, including sea urchin endomesoderm (Peter and Davidson, 2017), *C. elegans* gut development (Maduro, 2015) and secondary cell wall synthesis in vascular plants (Zhong and Ye, 2014). Coherent feedforward loops have multiple potential functions. One potential role is to introduce a deliberate delay into a developmental program (Mangan et al., 2003), though that seems unlikely in this case given the extremely rapid pace of *Ciona* fate specification. Another potential role is as persistence detectors and noise filters to prevent stochastic noise in transcription and/or signaling from inadvertently triggering inadvertent and irrevocable cell fate decisions (Peter and Davidson, 2017). This role seems more likely here, especially given the extreme stereotypy of ascidian cell fate specification. A similar feedforward loop involving FGF signaling has recently been identified in the *Ciona* cardiopharyngeal mesoderm (Razy-Krajka et al., 2018). FGF signaling plays a key role in numerous *Ciona* cell fate decisions, and more work will be needed to test the ubiquity of feedforward network motifs across different FGF-dependent cell types and more explicitly test the persistence detector/noise filter hypothesis.

The master regulator concept is used inconsistently in the literature, but it typically refers to a transcription factor that sits alone atop the regulatory cascade for the differentiation of a particular cell type that is not only necessary for the development of that cell type but also sufficient to transform other cell types to that fate when ectopically expressed (Chan and Kyba, 2013). Many putative master regulator TFs have been identified, including eyeless in Drosophila retinal cells (Gehring, 1996), MyoD for myoblasts (Tapscott et al., 1988; Weintraub et al., 1989) and a variety of TFs each controlling a specific CD4+ T cell lineage (Oestreich and Weinmann, 2012). More recently, however, there has been a growing recognition that these developmental cell fate decisions typically involve more complex regulatory networks with multiple essential regulators connected via feedback and/or feedforward loops (Davis and Rebay, 2017; Desplan, 1997; Kassar-Duchossoy et al., 2004; Wang et al., 2015). In *Ciona*, Bra was originally described as the master regulator of notochord fate (Hotta et al., 1999; Imai et al., 2006; Takahashi et al., 1999) due to its expression during notochord fate specification and its critical role in regulating notochord target genes inferred from both loss of function and gain of function experiments. The later description of several notochord genes (José-Edwards et al., 2011; Kugler et al., 2008) whose expression was Bra-independent suggested that this model might not be definitive. Examining a more comprehensive notochord expressed gene set, we have found that Bra alone is neither necessary nor sufficient to control expression of all notochord enriched genes. Based upon ectopic expression studies, CRISPR gene disruption and cis-regulatory analysis, we suggest a modified notochord GRN in which Bra and Foxa.a act through a feedforward loop in which neither acts as a canonical unitary master regulator (Figure 7). This is consistent with the observation in mouse embryos that although FoxA is an obligate upstream inducer of Bra expression (Ang and Rossant, 1994), it also has a temporally separable role later in notochord morphogenesis (Yamanaka et al., 2007). Bra and FoxA are coexpressed in axial tissues even in basal metazoans, so it is possible that this feedforward network may be evolutionarily ancient (Fritzenwanker et al., 2004).

Decades of research in developmental genetics support the idea that individual cell fate decisions are dominated by a small number of key transcription factors. The master regulator concept in which characteristic cell-type specific transcription factors are both necessary and sufficient to establish distinct cell states is appealing, as it is straightforward to understand and has important implications for the evolvability of body plans via conceptually simple cis-regulatory mutations in tissue-specific master regulators. The master regulator concept does not stand up to close inspection, however, as shown here for *Ciona* Bra and shown previously for other genes widely discussed as master regulators. We speculate that feed-forward interactions between small but non-singular sets of regulatory TFs may be characteristic features of cell fate decisions allowing a balance between the evolvability of body plans and the robustness of development to stochastic transcriptional noise.

## Materials and Methods

### *Ciona* husbandry and embryology

*Ciona robusta* (formerly known as *Ciona intestinalis* type A) (Pennati et al., 2015)were collected in San Diego and shipped to KSU by Marine Research and Educational Products Inc. (M-REP, San Diego, CA). Adult *Ciona* were maintained in a recirculating aquarium. Standard fertilization, dechorionation and electroporation protocols were used (Veeman et al., 2011). Staging is based upon the series of (Hotta et al., 2007).

### Electroporations and RNA prep

Fertilized dechorionated eggs were electroporated with either 40 ug of enhancer>Venus-Bra plasmid or 40 ug of control plasmid (*Bra*>GFP). The Foxa.a enhancer was tested separately from the Etr1 and Tbx6-r.b enhancers. For each of the three replicates, 400 experimental and 400 control embryos were collected at stage 19.5 into RNA lysis buffer and stored at - 80C until RNA purification. RNA was purified with the Zymogen Quick-RNA miniprep kit (R1054), genomic DNA was removed using the kit’s in column DNAse treatment step and purified RNA was eluted in 50 ul water, with a concentration of 52-60 ng/ul. Libraries were constructed with the TruSeq stranded mRNA library kit (Illumina) using standard protocols. Libraries were quality checked with an Agilent Tapestation and quantified by qPCR. Single end (1×100) sequencing was performed on an Illumina HiSeq 2500 at the KU Genome Sequencing Core (GSC) with a read depth of 33.5 to 38.5 million reads per sample for Foxa.a>Venus-Bra and its matched control or 13.5 to 15.4 million reads per sample for Etr1>Venus-Bra, Tbx6-r.b >Venus-Bra and their matched control.

For transcription factor combinations in the Etr1> plasmids, 33 ug of each plasmid was used per electroporation. The control plasmid was Etr1>H2B-Venus. Embryo collection and RNA preparation was as described above. Libraries were constructed with the NEBNext UltraII stranded mRNA library kit (NEB) using standard protocols and quality checks as above. Rapid Read single end (1×75) sequencing was performed on an Illumina NextSeq 550 at the KU GSC with a read depth of 26.6 to 39.6 million reads per sample.

### CRISPR-RNP injection and RNAseq

The guide RNA sequence used for Bra CRISPR was selected using scores calculated by CRISPOR (http://crispor.tefor.net) (Haeussler et al., 2016) and IDT (Integrated DNA Technologies, Inc. IA, USA). For GFP CRISPR, the sequence was obtained from Addgene (51762, https://www.addgene.org/)(Shalem et al., 2014). The sequences were TAAGCACATGACGAAACATG for Bra CRISPR and GGCCACAAGTTCAGCGTGTC for GFP CRISPR (PAM sequence not shown).

crRNAs, tracrRNAs and Cas9 protein were purchased from IDT (Alt-R® CRISPR-Cas9 crRNA, Alt-R® CRISPR-Cas9 tracrRNA and Alt-R® S.p. Cas9 Nuclease 3NLS or V3). The RNP mix were prepared just before injection by combining cr/tracrRNA, Cas9 protein and centrifuged HEPES/KCl buffer at final concentrations of 15 µM, 30 µM or 15 µM and 10 mM/75 mM, respectively, and then activated at 37°C for 5 min. RNP mix was injected into unfertilized eggs according to standard procedure (Yasuo and McDougall, 2018). Approximately 50 to 85 normal developing crispant embryos or control embryos were collected at early neurula (St. 14) and fixed by adding 350 µl of buffer RLT Plus, included in AllPrep DNA/RNA Mini Kit (Cat No./ID: 80204, QIAGEN, QIAGEN GmbH, Germany). The solution was then split in half before extracting genomic DNA and total RNA using the manufacturer’s protocol.

To evaluate the CRISPR mutagenesis efficiency, the Bra genomic locus was amplified (forward primer: TACGGCGCACTTTCAACAAA; reverse primer TTTGGTAGGGCGGGGTAATT) and Sanger sequenced from two directions (forward sequence primer GAGTGTGATTTGGAGGCAGA; reverse sequence primer ATGTGGATCCTGGGTTCGTA). The results were compared with ones from CRISPR GFP embryos using TIDE (https://tide.deskgen.com) (Brinkman et al., 2014) with default settings. The lower value for each sample (sequenced from F or R primer) was used as the samples TIDE score. Three replicates with high TIDE scores (∼70%) were used for constructing RNA libraries. RNA libraries were constructed, tested and sequenced as described for the Bra/Mnx/Foxa.a ectopic expression experiment. Read depth was 19.2 to 23.4 million reads per sample.

### Calculation of theoretical maximum decrease by Bra CRISPR

We previously showed using RNAseq on FACS-sorted notochord and not-notochord cells that there is a smooth distribution of notochord enrichment values and not a bimodal distribution of ‘notochord-specific’ vs ‘notochord-enriched’ genes. In our Brachyury CRISPR experiments, we assume that knocking out Brachyury will affect gene expression in notochord cells but not in other cell types. The fold change observed by whole-embryo RNAseq for a given gene is thus a function of not only the degree to which that gene is dependent on Brachyury but also the degree to which it is enriched in the notochord. Since the total expression of a notochord-enriched gene in whole embryo RNAseq experiments is equal to total notochord expression + total non-notochord expression, then total expression = aN + (1-a)E, where N is equal to expression in notochord cells, E is the expression in non-notochord cells and a is the fraction of the embryo comprised of notochord cells. Fold perturbation effect (Fp) is equal to total expression in Bra crispant embryos/total expression in control embryos. The maximum fold change of perturbation expected in a Bra CRISPR experiment using whole embryo RNAseq will be if the notochord expression is eliminated leaving only the background expression outside the notochord:

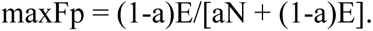

We have previously estimated notochord fold enrichment (Fe) by FACS-Seq, with Fe = N/E or N = Fe*E. Therefore maxFp = (1-a)E/[a*Fe*E + (1-a)E] which can be simplified to (1-a)/(A*Fe + 1 –a). All 10 notochord founder cells are specified by the 112-cell stage, so we estimate that 10/112 (∼9%) of the embryo by mass at later stages consists of notochord cells:

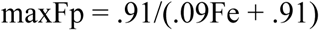

### Drug treatments

For the RNAseq experiment, U0126 was dissolved in DMSO at 4 mM and used at a final concentration of 4 micromolar. DMSO alone (at 1:1000 dilution) was used as a control. Embryos were treated at the 32 cell stage, then collected at the 64 cell stage and 112 cell stage. RNA purification, library preparation and sequencing was as described for Bra/Mnx/Foxa.a ectopic expression experiments. Read depth was 18.7 to 23 million reads per sample.

For in situ hybridization, embryos were treated with 10 micromolar Cytochalasin B at the 112 cell stage after U0126/DMSO treatment at the 32 cell stage, then fixed at stage 16 as described in (Reeves et al., 2014)

### RNAseq analysis and bioinformatics

Tophat (Trapnell et al., 2013, 2012) was used to align reads to the *Ciona* KH2012 gene models (KH.KHGene.2012.gff3, retrieved from ANISEED, http://www.aniseed.cnrs.fr/) (Brozovic et al., 2016; Dardaillon et al., 2020). The standard DEseq pipeline (Love et al., 2014) was used to calculate read counts and test for differential expression. Statistical significance was reported as adjusted p-values.

### TFBS enrichment analysis

We used bedtools (Quinlan and Hall, 2010) and custom scripts to extract putative regulatory regions near the predicted transcriptional start sites of both notochord-enriched and control gene sets. We performed separate analyses using our FACS RNAseq data from both stage 16 and stage 19. For each timepoint, the notochord-enriched gene set included all genes enriched at least two-fold in notochord. As control sequences, we used the 1000 most notochord depleted genes at each stage. See supplemental materials for code details. Putative regulatory regions were defined as DNA sequences 1500 bp upstream and downstream of transcription start sites. For genes predicted to be expressed in operons, we used the operon TSS. We then subtracted coding exon sequences and removed any features less than 100 bps. We removed any sequences from analysis that were present in both sets at a given stage. TFBS enrichment was calculated using the 2018 JASPAR vertebrate TFBS profile set (JASPAR; http://jaspar.genereg.net)(Khan et al., 2018) and the oPOSSUM 3.0 Single-Site Analysis algorithm (Kwon et al., 2012) with default settings. The Z-scores (which compare the rate of occurrence of a given TFBS in the test set vs the control set) are reported.

### Analysis of mutated Bra enhancer constructs

The Bra reporter TFBS mutation analysis made use of a 377 bp fragment matched to an ATAC-seq peak over the upstream region and first exon of the Bra locus (KhS1404:5981-6357), cloned into a vector containing a minimal promoter from *Ciona* Friend of Gata (bpFOG) and a Venus YFP reporter (pX2+bpFOG>UNC76:Venus (Stolfi et al., 2015)). Predicted transcription factor binding sites were identified using JASPAR 2018 PWMs and the FIMO scanner(http://meme-suite.org/doc/fimo.html)(Grant et al., 2011) using a p-value threshold of 0.001. Mutant sequences designed to disrupt specific binding sites were rescanned by FIMO to ensure that they did not affect nearby sites or inadvertently introduce new sites. Control and mutant sequences were all synthesized as gBlocks and verified by Sanger sequencing after Gibson cloning into the reporter vector. All constructs were electroporated in two (Zic mutant) or three (Ets and Fox mutants) different biological replicates in different combinations, at 50 micrograms each. All experiments included the wildtype control plasmid as one of the electroporated constructs. For each experiment, embryos were fixed at stage 21, immunostained for reporter expression, and cleared in Murray’s Clear.

At least 10 embryos were imaged per construct per replicate using a 40x 1.3NA objective on a Zeiss 880 confocal microscope using uniform imaging settings. These were randomly selected without inspecting reporter staining from embryos showing normal embryonic morphology by phalloidin staining. 3D image stacks were used to reconstruct a computationally straightened slice along the length of the notochord (described in (Harder et al., 2019)) and a thick line trace along the notochord was summed to quantify a reporter expression value for each embryo. Expression of electroporated transgenes in *Ciona* is mosaic, so expression at the level of individual embryos is quite variable. This represents both embryo to embryo variation in the number of expressing cells and how brightly they are expressing, as well as variation in transfection efficiency between different electroporations. We controlled for this by imaging multiple randomly chosen embryos for each replicate and electroporating multiple replicates for each construct as described above. The data were normalized at the level of the entire dataset to give the control plasmid a median log-transformed value of 1.

### In situ hybridization and probes

Probe synthesis, embryo collection, in situ hybridization and imaging were performed as described in (Reeves et al., 2014). Probes for Bra (Reeves et al., 2014),C7.633 and C1.1281 (Reeves et al., 2017) were previously described. Probe sequences for C11.313, C3.815, C2.866, C2.33, C9.698, C7.260 and L170.42 were amplified from cDNA, then cloned into pBSII SK(-). Primers used are provided in Supp Table 6.

### Plasmid construction details

Foxa.a>Bra-Venus was previously described (Reeves et al., 2017; Takahashi et al., 1999). For Etr1>Bra-Venus, the Foxa.a enhancer was removed by XhoI/XbaI digest, and replaced with PCR amplified enhancers for Etr1 (forward primer TCGACCACGGAGTTAATTG; reverse primer TCTGGATAAAGCAATACATACGAG) and Tbx6-r.b (forward primer AATGTAGCGTCGCTTCAC; reverse primer GGAATCATATTCGCCATAGTC) by Gibson cloning (NEB kit). For Etr1>Mnx-HA, full length Mnx coding sequence (PCR amplified from genomic DNA) was Gibson cloned upstream of the HA tag in SwaI-Rfa-HA, replacing the Gateway cloning cassette, and then the Etr enhancer was cloned into XhoI/XbaI as above. For Etr1>Foxa.a-Myc we used three fragment Gibson cloning to combine the Etr enhancer (PCR amplified from Etr1<Bra-Venus), the Foxa.a coding sequence (PCR amplified from genomic DNA with myc tag added in primer) and the plasmid backbone of SwaI-Venus-Rfa (ref) outside of the Venus-Rfa region.

## Supporting information

Script for extracting putative regulatory regions for TFBS enrichment analysis

## Funding and Acknowledgements

This work was supported by NIH award 1R01HD085909 to MV. We thank Julie Hix for technical assistance. We also acknowledge support from the KSU CVM confocal core, the KSU Beocat cluster computing core, the KSU Integrated Genomics Facility, and the KU CMADP COBRE Genome Sequencing core.

